# The Effects of Alcohol and Cannabinoid Exposure during the Brain Growth Spurt on Behavioral Development in Rats

**DOI:** 10.1101/513796

**Authors:** Kristen R. Breit, Brandonn Zamudio, Jennifer D. Thomas

## Abstract

Cannabis is the most commonly used illicit drug among pregnant women. Moreover, over half of pregnant women who are consuming cannabis are also consuming alcohol; however, the consequences of combined prenatal alcohol and cannabis exposure on fetal development are not well understood. The current study examined behavioral development following exposure to ethanol (EtOH) and/or CP-55,940 (CP), a cannabinoid receptor agonist. From postnatal days (PD) 4-9, a period of brain development equivalent to the third trimester, Sprague-Dawley rats received EtOH (5.25g/kg/day) or sham intubation, as well as CP (0.4 mg/kg/day) or vehicle. All subjects were tested on open field activity (PD 18-21), spatial learning (PD 40-46), and elevated plus maze (PD 30) tasks. Both EtOH and CP increased locomotor activity in the open field, and the combination produced more severe overactivity than either exposure alone. Similarly, increases in thigmotaxis in the Morris water maze were caused by either EtOH or CP alone, and were more severe with combined exposure, although only EtOH impaired spatial learning. Finally, developmental CP, but not EtOH, increased time spent in the open arms on the elevated plus maze. Overall, these data indicate that EtOH and CP produce some independent domain-specific effects, but many effects of EtOH and CP on behavior were additive. Importantly, these data suggest that combined prenatal exposure to alcohol and cannabis may be more damaging to the developing fetus, which has implications for the lives of affected individuals and families and also for establishing public health policy.

## 1. Introduction

The adverse consequences of prenatal alcohol exposure have been extensively studied over the past 40 years (Riley et al., 2011). Individuals exposed to alcohol prenatally may suffer from a range of physical, neurological, and behavioral consequences referred to as fetal alcohol spectrum disorders (FASD). FASD include growth deficits, facial dysmorphology, as well as altered cognitive, emotional, and behavioral functions (Riley et al., 2011). Approximately 1 in 10 women in the U.S. report some alcohol consumption during pregnancy (Centers for Disease Control and Prevention, 2015; Substance Use and Mental Health Services Administration, 2014). Thus, FASD continues to pose a serious public health problem in the U.S., as well as around the globe (Roozen et al., 2016).

Prenatal alcohol exposure alters development of numerous brain regions, leading to structural and functional alterations in such areas as the basal ganglia, cerebellum, cortex and hippocampus (Moore et al., 2014) as well as white matter tracts (Taylor et al., 2015; Uban et al., 2017). Alcohol-induced neuropathology can lead to behavioral alterations and impairments in learning, attention, executive functioning, and emotional regulation, all of which can cause serious problems in school and daily life (Khoury et al., 2015; Mattson et al., 2011; Norman et al., 2013). Similarly, animal studies have also shown that developmental alcohol exposure alters brain development, producing a variety of behavioral alterations, including hyperactivity, increased anxiety-related behaviors, and impaired learning and memory (Schneider et al., 2011).

However, alcohol is not the only drug of abuse consumed by pregnant women. Recent reports show that 5.4% of pregnant women reported using illicit drugs (Substance Use and Mental Health Services Administration, 2014), with higher rates of use among younger pregnant women (15-17 years; 14.6%) (Substance Use and Mental Health Services Administration, 2014). The most commonly used illicit drug among women of reproductive age is cannabis (Substance Use and Mental Health Services Administration, 2014), with prevalence rates of 3-4% among pregnant women in Western countries (Ebrahim et al., 2003; El Marroun et al., 2011). Similar to alcohol, the primary psychoactive constituent in cannabis (∆9-tetrahydrocannabinol [THC]) and its metabolites can freely cross the placenta and directly affect the fetus (Gómez et al., 2003). Of particular concern are the steadily increasing amounts of THC in cannabis products. In 2016, the average potency of THC in cannabis-related products was 11% (Botticelli, 2017); this number has continually risen from 3.4% in 1993 and is even higher among synthetic variations (Botticelli, 2016; Mehmedic et al., 2010). Importantly, many women perceive cannabis as safe to use during pregnancy (Saint Louis, 2017).

Although the dangers of alcohol consumption during pregnancy are well established, much less is known of the consequences of prenatal cannabis exposure, particularly at the high levels consumed today. There are few prospective clinical studies (such as Generation R) (Jaddoe et al., 2012) examining the effects of prenatal cannabis exposure (Huizink, 2014), and results from these and retrospective studies are mixed, likely due to differences in cannabis exposure levels, prospective vs. retrospective approaches, confounding of other drug use like tobacco, age and nature of outcome measures, as well as a host of other methodological, maternal, and environmental factors. In general, evidence suggests that prenatal cannabis exposure may alter emotional, behavioral, and cognitive development in clinical populations, particularly in executive functioning domains (Huizink, 2014). Unfortunately, the consequences of prenatal exposure to the increased THC potencies available today will not be known for years to come. Animal studies are similarly inconsistent; while prenatal THC exposure may not cause neuronal cell death in areas such as the hippocampus, cortex, and thalamus (Hansen et al., 2008), animal model studies do illustrate that prenatal cannabinoid exposure can produce hyperactivity (Huizink et al., 2006), increase anxiety-related behaviors (Goldschmidt et al., 2004), and impair working (Smith et al., 2006) and long-term memory (Mereu et al., 2003), although behavioral results are mixed. In addition, perinatal cannabinoid exposure has been shown to impair reproductive functioning in male mice (D’Alterio et al., 1979) and disrupts neurodevelopmental CB_1_ signaling later in adulthood (de Salas-Quiroga et al., 2015). Similar to clinical studies, mixed results likely vary based on differences in timing, dose and form of cannabinoid, outcome measures, and nature of control groups (Abel et al., 1986; Huizink, 2014; Schneider, 2009; Trezza et al., 2012).

Moreover, the effects of concurrent exposure to alcohol and cannabis are not known, despite the high rates of co-use reported among women of child-bearing age. A recent analysis found that 8.7% of females use both alcohol and cannabis, with 5.5% reporting simultaneous use, suggesting that individuals who use both drugs tend to consume them at the same time (Subbaraman et al., 2015). Higher rates of simultaneous alcohol and cannabis use (15.3%) were observed in the 18-29 year age group (Subbaraman et al., 2015), which is consistent with other recent data showing that 19.5% of individuals under age 21 reported smoking cannabis within 2 hours of consuming alcohol (Substance Use and Mental Health Services Administration, 2014).

Importantly, cannabis is also the illicit drug most commonly used simultaneously with alcohol among women who binge drink during pregnancy (Bhuvaneswar et al., 2007); approximately 50% of pregnant women who have reported consuming cannabis were also drinking alcohol (Substance Abuse and Mental Health Services Administration, 2015). These numbers may be even higher than reports suggest; accurate data reflecting concurrent use and exposure levels among pregnant women are difficult to obtain, not only because women who consume either alcohol or cannabis frequently under-report their usage (Lange et al., 2014; Lendoiro et al., 2013; Midanik, 1988), but also because of the rapidly changing legalization, accessibility, and potency of cannabis. In addition, approximately 50% of pregnancies are unplanned. Thus, fetuses may be exposed to these drugs before pregnancy is recognized, which is alarming given the high rates of concurrent use.

The possibility that these drugs may interact with one another is strengthened by the role of endogenous cannabinoids in alcohol-induced neurodegenerative pathways (Subbanna et al., 2013), depression of synaptic activity (Nagre et al., 2015), and impaired DNA methylation (Basavarajappa et al., 2008). Furthermore, alcohol can alter the development of endocannabinoid systems, including neuronal communication and circuit formation, which may lead to deficits in neuronal plasticity (Basavarajappa, 2015). Thus, concurrent exposure during early development may be even more detrimental to the developing fetus than either drug alone.

However, there is limited research on co-exposure to prenatal alcohol and cannabis on brain and behavioral development. Most clinical (Fried et al., 1987; Fried et al., 1992; Fried et al., 1990; Richardson et al., 2002) and animal studies (Hansen et al., 2008; Subbanna et al., 2013) have focused on the effects of each drug separately (Fried et al., 1987; Fried et al., 1992; Fried et al., 1990; Richardson et al., 2002), rather than the combination of effects. Research that has examined the combined effects suggest that concurrent cannabinoid and alcohol exposure can significantly increase fetal toxicity (Abel et al., 1987) and synergistically increase neurotoxicity in the hippocampus (Hansen et al., 2008) in rodents. In addition, exposure to cannabinoids on gestational day (GD) 8 produces craniological, ocular, and brain abnormalities in a dose-dependent manner (Gilbert et al., 2016), whereas concurrent alcohol and cannabinoid exposure yielded synergistic effects, showing greater physical abnormalities than exposure to either alcohol or cannabinoids alone (Fish et al., 2017; Fish et al., 2016). Nonetheless, research on behavioral development following co-exposure is lacking. Limited previous clinical (Goldschmidt et al., 2004) and animal studies (Abel et al., 1990) have failed to find any interactions of prenatal alcohol and THC on behavioral development, but these studies investigated low doses of THC or alcohol, notably lower than the THC levels consumed by individuals today.

To examine the possible consequences of combined developmental exposure to alcohol and cannabinoids on behavioral development, we used a rat model of drug exposure during the 3^rd^ trimester brain growth spurt equivalent (postnatal days (PD) 4-9). Importantly, this is a period of development particularly sensitive to ethanol (Goodlett et al., 1991; Olney et al., 2002) and a period in which the endogenous cannabinoid system plays an important role in neuronal development (Fernández-Ruiz et al., 2000). The brain growth spurt is characterized by neuronal maturation, including axonal growth, dendritic arborization, as well as high rates of synaptogenesis, gliogenesis, and myelination (Dobbing et al., 1979; Gauda, 2006). Additionally, CB_1_ receptor levels rapidly increase in numerous brain regions during this time (Belue et al., 1995; Berrendero et al., 1999; Rodriguez de Fonseca et al., 1993).

The current study used the synthetic CB_1_ and CB_2_ receptor agonist CP-55,940 (CP) to mimic the effects of THC. CP is as an ingredient in synthetic marijuana preparations (Berkovitz et al., 2011) and is commonly used in cannabinoid research (Tai et al., 2014) as it has a similar duration of action, peak effect, and neurobehavioral effects as THC (McGregor et al., 1996). CP was administered in a manner equivalent to a moderate-high dose in humans (Desrosiers et al., 2014; García-Gil et al., 1999; Little et al., 1988) and combined with a well-established binge-like dose of alcohol (McGough et al., 2009; Thomas et al., 2008; Thomas et al., 2012).

All subjects were tested on a battery of behavioral tests to investigate effects of combined exposure to alcohol and cannabinoids on behavioral development. The behavioral battery included an open-field activity chamber (activity levels), an elevated plus maze (anxiety-related behaviors), and a Morris water maze task (visuospatial learning). Following the behavioral battery, gross brain weights (g) were measured.

## 2. Material and Methods

The goal of the current study was to examine the effects of combined neonatal alcohol and cannabinoid exposure on behavioral development (see Figure 1). All procedures and behavioral paradigms included in this study were approved by the San Diego State University (SDSU) Institutional Animal Care and Use Committee (IACUC) and are in accordance with the National Institute of Health’s *Guide for the Care and Use of Laboratory Animals*.

**Figure 1.**
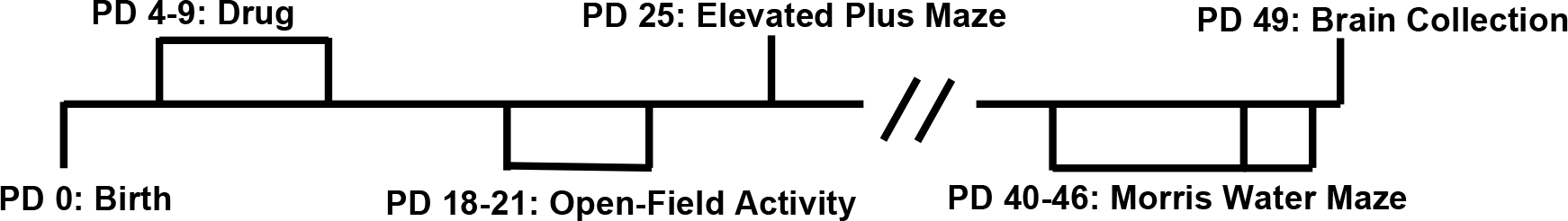
Timeline of drug exposure, behavioral testing, and tissue collection.

### 2.1. Subjects

For breeding, one male and one female Sprague-Dawley rat were housed together overnight at the SDSU Animal Care facilities at the Center for Behavioral Teratology. Once a seminal plug was present, dams were individually housed in standard plastic cages (gestational [GD] 0) and left undisturbed until the day of delivery (typically GD 22), except for routine husbandry. On postnatal day (PD) 1, the day after birth, litters were pseudo-randomly culled to 8 pups with equal numbers of male and female subjects (whenever possible). Subjects were then randomly assigned to each exposure group; no more than one sex pair per litter was used within each exposure condition to control for potential litter effects. A total of 142 subjects were treated from 18 litters.

### 2.2. Developmental Alcohol and Cannabinoid Exposure

From postnatal days (PD) 4-9, a time period equivalent to the human third trimester “brain growth spurt,” half of the subjects were intragastrically intubated with ethanol (EtOH, 5.25 g/kg/day) dissolved in an artificial milk diet (11.9% v/v, 27.5 mL/kg, twice per day, 2 hours apart), followed by 2 additional feedings of milk diet only (Goodlett et al., 1997). Briefly, the intubation procedure involved lubricating a thin, silastic tubing (PE-50) with corn oil, inserting the tubing into the pup’s mouth to be swallowed into the stomach, and injecting the milk diet through the tubing (Goodlett et al., 1997). The other half of subjects received sham intubations, which involved full insertion of the tubing, but with no EtOH or milk exposure. In addition, all subjects were injected (i.p.10 ml/kg) with either CP (0.4 mg/kg/day) or the vehicle (VEH; 10% dimethylsulfoxide [DMSO] and sterilized saline). Injections utilized a 30-gauge needle with a guide to allow only the tip of the needle to enter the i.p. area. All pups were removed from the dam simultaneously and maintained on a heating pad (110 °F) throughout the exposure period. Each intubation and injection took approximately one minute per pup to complete; litters were returned to the dam within 10 minutes of separation.

Offspring were tattooed for identification purposes with non-toxic veterinary tattoo ink on PD 7, allowing the experimenter to remain blind during later behavioral testing; prior to PD 7, a non-toxic marker was used for identification. Subjects were weaned on PD 21 and group-housed by sex on PD 28. All subjects were housed at a constant humidity and temperature (21±1°C) in plastic cages with woodchips, and exposed to a 12-hour light/dark cycle, receiving food and water *ad libitum*.

#### 2.2.1. Drug Preparation

Ethanol (11.9% v/v) was added to an artificial milk diet (West et al., 1984). CP-55,940 (CP; Enzo Life Sciences, NY) was dissolved into a stock solution (5 mg of CP dissolved into 2mL of 100% Dimethyl Sulfoxide [DMSO] Sigma-Aldrich, MO) and kept at −20°C until daily injections volumes were made. Daily injection volumes were prepared by combining the CP stock solution with the vehicle (10% DMSO in physiological saline) to the appropriate final dose (0.4 mg/kg/day).

#### 2.2.2. Behavioral Testing

##### 2.2.2.1 Activity Levels

From PD 18-21, activity levels in an open field were measured in all subjects during the dark cycle (evening). For 60 min, all subjects placed in separate, dark open-field activity chambers (41 × 46 × 38 cm; Hamilton-Kinder) equipped with fans for ventilation and white noise. Prior to weaning (PD 21), subjects were removed from the dam simultaneously before being placed in the chambers (a potential stressor), while on PD 21 weaning had already occurred. Activity levels were recorded in 5-min bins via interruptions of infrared beams, indicating overall activity levels, location and time spent in different areas of the chamber, and behaviors such as rearing.

##### 2.2.2.2. Anxiety-Related Behaviors

On PD 25, anxiety-related behaviors were measured in all subjects using an elevated plus maze test. The plus sign-shaped maze is elevated 50 cm above the floor, with two exposed arms and two enclosed arms (50 × 10 cm); the maze is made of clear Plexiglas except for the black sides of the enclosed arms (40 cm). Subjects were placed in the center of the maze and allowed to roam freely for 5 min. A video camera recorded subjects’ locations on the maze (open vs closed arms), time spent in each area, as well as grooming and rearing behaviors, which were later coded using OdLog software. The center of the maze was coded as an open arm.

##### 2.2.2.3. Visuospatial Learning and Memory

From PD 40-46, all subjects were tested for visuospatial memory performance using a Morris water maze. Subjects were required to use spatial cues in the room to remember the location of a clear Plexiglas platform (4-inch diameter) hidden 1 inch below the water surface of a black tank (48-inch diameter) filled with water. All data were recorded via a video tracking system interfaced with Water2020 software (HVS Image).

For acquisition, subjects were tested for 4 trials per day over 6 consecutive days with an inter-trial interval of 3-5 min. For each trial, the platform location remained consistent, while the starting position changed. Subjects were given 60 sec to find and swim to the platform; if subjects failed to find the platform, they were guided. Once on the platform, subjects were given 10 additional sec to observe spatial location. Distance and latency traveled to find the platform, as well as heading angle, and thigmotaxis (“wall-hugging,” outer 10% of the pool) were measured. Twenty-four hours after the last acquisition trial, subjects were given a 60-sec probe trial, in which the platform was removed and memory of the platform location was measured by examining time spent and passes through the platform area.

#### 2.2.3. Brain Collection

Brains were collected from all subjects via intracardial perfusion between PD 47-51. Subjects were given a lethal dose of pentobarbital (10% pentobarbital in sterile saline, 3 mL/kg, i.p.) and perfused with a 4% paraformaldehyde solution. Once perfused, the forebrain and cerebellum were collected and weighed.

### 2.3. Statistical Analyses

All data were analyzed using the Statistical Package for the Social Sciences (SPSS, version 24). All analyses used a 2 (EtOH exposure: EtOH, Sham) × 2 (CP exposure: CP, VEH) × 2 (sex: female, male) ANOVA. Repeated measures ANOVAs for Day, 5-minute Time Bin (Bin), and/or Trial were used when applicable. Post hoc analyses were conducted using Student-Newman-Keuls. Means (M) and standard errors of the mean (SEM) are reported when applicable. All significance levels were set at *p* < 0.05. In addition, to determine if CP had effects by itself or altered ethanol’s effects, specific post-hoc comparisons were made between CP and controls and between CP+EtOH and EtOH groups.

## 3. Results

At least 10 subjects per exposure group and sex completed all behavioral tasks (EtOH + CP: 23 [13 F, 10 M], EtOH + VEH: 30 [16 F, 14 M], Sham + CP: 30 [15 F, 15 M], Sham + VEH: 28 [13 F, 15 M]).

### 3.1. Body Weights

On the first day of the open-field activity testing (PD 18), subjects exposed to EtOH during development weighed significantly less than sham-intubated subjects (F[1,103] = 38.19, *p* < 0.001; Figure 2). There was also a main effect of CP (F[1,103] = 7.40, *p* < 0.01); although there was no significant interaction of EtOH*CP on body weights during this period, the main effect of CP was driven by reductions in body weight among subjects exposed to EtOH+CP. Although there were no effects of sex on body weight during open-field activity, males weighed more than females during the remaining behavioral tests (*p’*s < 0.001). Sex did not interact with either developmental EtOH or CP exposure at any time.

**Figure 2.**
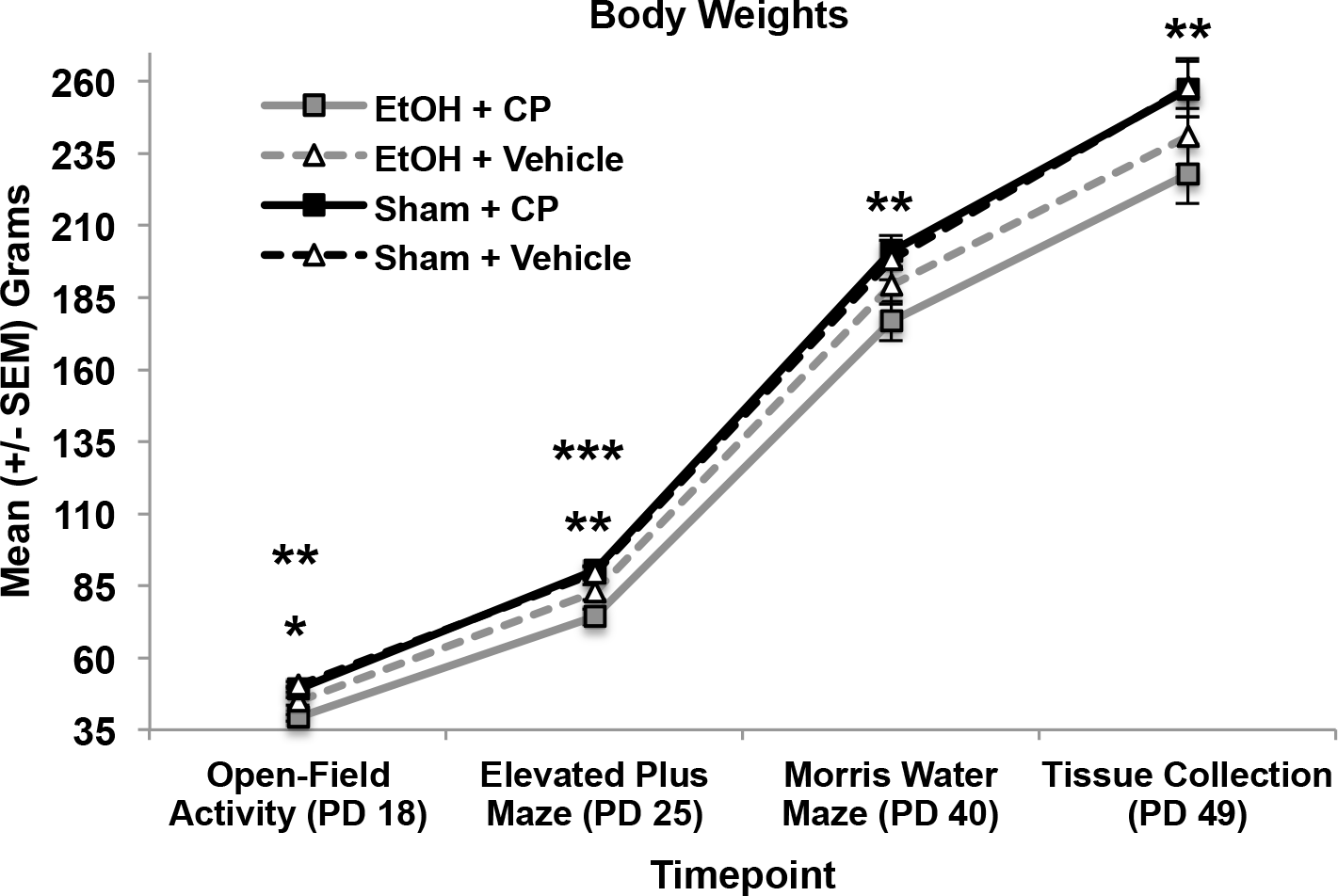
Developmental EtOH reduced body weight, an effect that persisted throughout early adulthood. CP exposure exacerbated the ethanol-related growth reductions up to PD 25. *** = EtOH+CP < EtOH+Vehicle; ** = EtOH < Sham; * = CP < Vehicle.

On PD 25, during the elevated plus maze task, there was an interaction of EtOH*CP exposure on body weights (F[1,103] = 6.26, *p* < 0.05). Subjects exposed to combined EtOH+CP during development weighed less than those exposed to EtOH alone (F[1,51] = 7.32, *p* < 0.01), whereas subjects exposed to only CP did not weigh significantly different from controls (F[1,56] = 0.15, *p* = 0.70). However, by the first day of the Morris water maze test (PD 40), only a main effect of developmental EtOH exposure persisted (F[1,103] = 8.45, *p* < 0.01), as EtOH-exposed subjects weighed less than their sham-intubated counterparts, regardless of CP exposure. EtOH-related reductions in body weight persisted until sacrifice (PD 49) (F[1,103] = 11.41, *p* < 0.01). In contrast, developmental CP exposure had no significant long-lasting effects on body weights.

### 3.2. Activity Levels

All activity measures were initially analyzed using a 4 (Day) × 12 (Bin) × 2 (EtOH) × 2 (CP) × 2 (Sex) repeated measures ANOVA. All subjects habituated within and across testing days on all activity measures, producing main effects of Day (*p’*s < 0.001), Bin (*p’*s < 0.001), and interaction of Day*Bin (*p’*s < 0.001). Habituation differences among treatment groups are noted, if applicable. No differences between male or female subjects were observed in any activity measure.

#### 3.2.1. Overall Activity

Subjects exposed to EtOH during development were more active than sham-intubated subjects, as evident by increased total distance traveled (F[1,103] = 28.9, *p* < 0.001; Figure 3A). In addition, EtOH-exposed subjects habituated slower, producing a Bin*EtOH interaction (F[11,1133] = 4.2, *p* < 0.001; Figure 3B). Although the interaction of EtOH and CP failed to reach statistical significance, CP by itself did significantly increase total distance traveled in sham, but not EtOH-exposed, subjects (F[1,56) = 5.1, *p* < 0.05; Figure 3A). In addition, locomotor activity of subjects exposed to CP did not habituate to the same level as controls, producing a Day*Bin*CP interaction (F[33,3399] = 1.5, *p* < 0.05). Similar effects were observed in the total number of beam breaks (data not shown).

**Figure 3.**
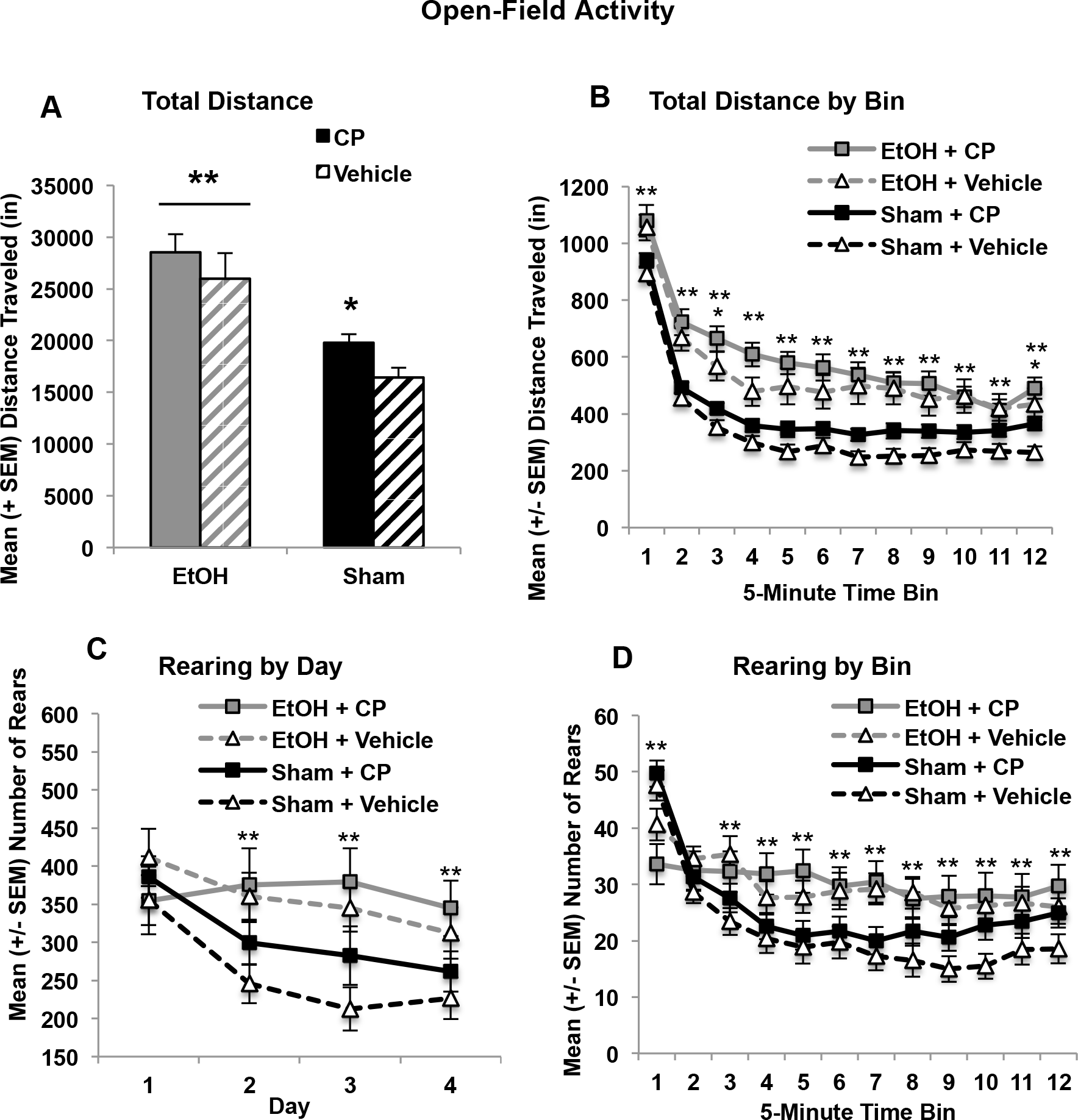
Ethanol exposure during the 3^rd^ trimester equivalent increased overall (summed) locomotor activity in the open-field chamber (A) and slowed the average habituation across days (B). Similarly, subjects exposed to ethanol showed less habituation of rearing by day (bins averaged) compared to their Sham counterparts (C). CP exposure also reduced habituation of total and average locomotor activity (A,B) and reduced habituation of rearing averaged across days (D). * = CP > Vehicle; ** = EtOH > Sham.

#### 3.2.2. Center-Related Activity

In contrast, when locomotor activity in the center of the chamber was examined, the combination of developmental EtOH+CP exposure produced more severe overactivity than either exposure by itself. Separately, both EtOH and CP increased locomotor activity in the center of the chamber, producing main effects of EtOH (F[1,103] = 53.1, *p* < 0.001) and CP (F[1,103] = 8.1, *p* < 0.01; Figure 4A). However, subjects exposed to combined EtOH+CP took significantly longer to habituate compared to all other groups (Figure 4B), producing a 3-way interaction of Bin*EtOH*CP (F[11,1133] = 1.9, *p* < 0.05). Importantly, although the combination of EtOH and CP increased locomotor activity in the center of the chamber, only developmental EtOH exposure significantly increased the time spent in the center of the chamber (F[1,103] = 16.4, *p* < 0.001; Figures 4C, 4D). CP exposure had no significant effect on time spent in the center, nor did ethanol and CP interact on this measure. Similar effects were seen with entries into the center of the chamber (data not shown).

**Figure 4.**
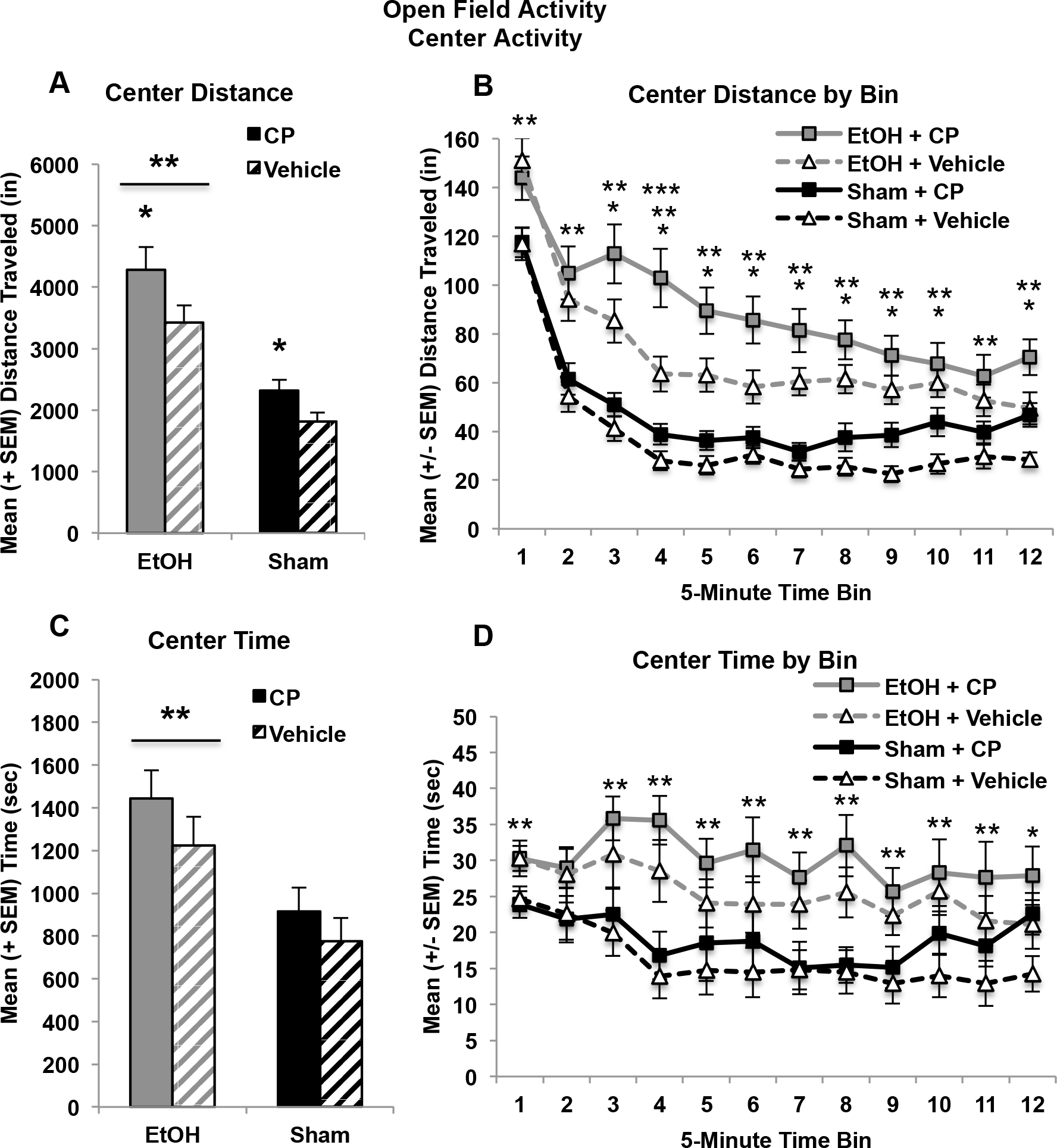
Both developmental EtOH and CP exposure separately increased the total locomotor activity in the center of the chamber (A), and the combination impaired habitation when averaged across days (B). However, only developmental EtOH exposure significantly increased the total and average time spent in the center of the chamber (C, D). * = CP > Vehicle; ** = EtOH > Sham; *** = CP × EtOH Interaction.

#### 3.2.3. Exploratory Activity

Overall, EtOH exposure increased rearing in the chamber (F[1,103] = 7.1, *p* < 0.01), regardless of CP exposure. A Day*EtOH interaction was present (F[3,309] = 5.0, *p* < 0.01), as EtOH-exposed subjects exhibited less habituation over days (Figure 3C), rearing significantly more during the last three days of open field activity testing (PD 19-21; *p’*s < 0.01). In addition, a three-way interaction between Day*Bin*CP was evident (F[33,3399] = 1.7, *p* < 0.01), as subjects exposed to CP took longer to habituate rearing across and within sessions (Figure 3D).

### 3.3. Anxiety-Related Behaviors

The elevated plus maze utilizes rats’ natural aversion to open spaces and other behaviors to infer levels of anxiety. All measures on this task were analyzed using a 2 (EtOH) × 2 (CP) × 2 (Sex) ANOVA. Exposure to EtOH during development did not significantly alter the time spent on the open versus closed arms of the elevated plus maze, nor did it alter the frequency of open, closed, or total arm entries. In contrast, developmental CP exposure increased the time spent on the open arms of the elevated plus maze (F[1,103] = 5.91, *p* < 0.05; Figure 5A). Although an interaction of EtOH*CP was not significant, it should be noted that this effect was driven by an increase in open arm time among subjects exposed only to CP compared to controls (F[1,56] = 9.39, *p* < 0.01); CP had no effect among EtOH-exposed subjects (F[1,51] = 0.43, *p* = 0.52).

Importantly, CP exposure did not alter the frequency (Figure 5B) or the ratio of different arm entry, suggesting that CP-exposed subjects specifically spent more time in the open areas of the maze.

**Figure 5.**
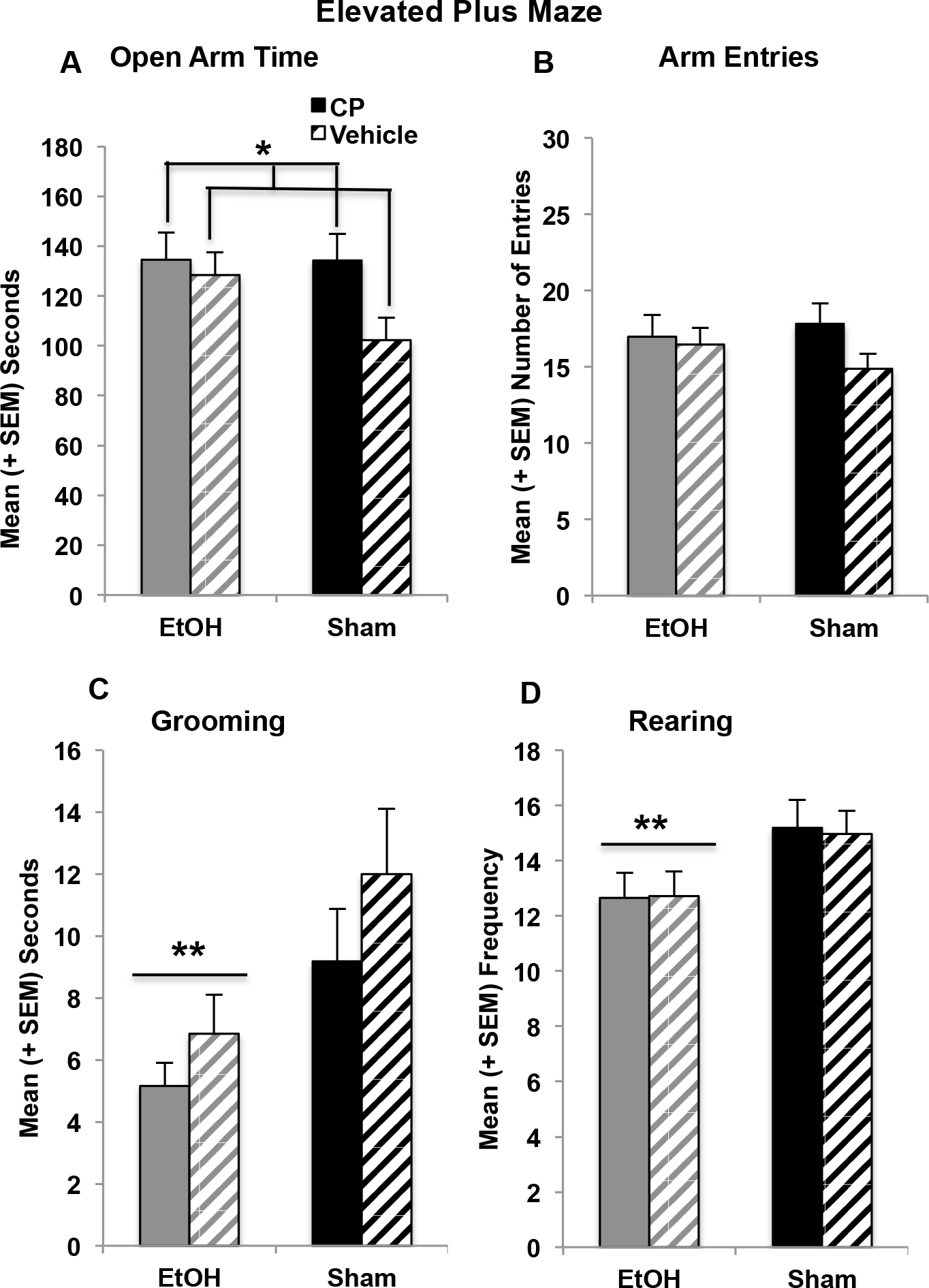
CP exposure during the 3^rd^ trimester equivalent increased the total time spent on the open arms of the elevated plus maze, driven by a CP-related increase among sham-intubated subjects (A). In contrast, EtOH exposure decreased the time spent grooming (C) and the number of rearing (D) behaviors on the maze. No groups differed in overall activity on the plus maze (B). * = CP > Vehicle; ** = EtOH < Sham.

However, EtOH exposure decreased the time spent grooming (F[1,103 = 9.39, *p* < 0.01; Figure 5C), grooming frequency (F[1,103] = 4.55, *p* < 0.05), and average grooming bout duration (F[1,103] = 4.98, *p* < 0.05). In addition, developmental EtOH exposure also reduced the frequency of rearing, an exploratory behavior (F[1,103] = 10.19. *p* < 0.01; Figure 5D), but not affect the total time spent rearing or average rearing bout. Neither grooming nor rearing behaviors were altered by developmental CP exposure. Moreover, there were no interactions between EtOH*CP or effects of Sex on any behaviors on the elevated plus maze.

### 3.4. Spatial Learning

#### 3.4.1. Acquisition

All acquisition data were initially analyzed using a 6 (Day) × 2 (EtOH) × 2 (CP) × 2 (Sex) repeated measures ANOVA. Although performance in all subjects improved over acquisition days (effects of day, *p’*s < 0.001), subjects exposed to EtOH during development exhibited impairments in spatial learning. EtOH-exposed subjects learned more slowly than controls and required longer path lengths to find the platform compared to controls, producing an interaction of Day*EtOH (F[5,515] = 2.19, *p* = 0.05), as well as a main effect of EtOH (F[1,103] = 39.90, *p* < 0.001; Figure 6A). Similar effects were seen with latency to find the platform, even though EtOH-exposed subjects did swim faster. In addition, EtOH-exposed subjects were less precise in aiming toward the platform, exhibiting larger heading angles (F[1,103] = 15.73, *p* < 0.001; Figure 6B). Finally, males performed better than females on all acquisition measures (*p’*s < 0.01).

**Figure 6.**
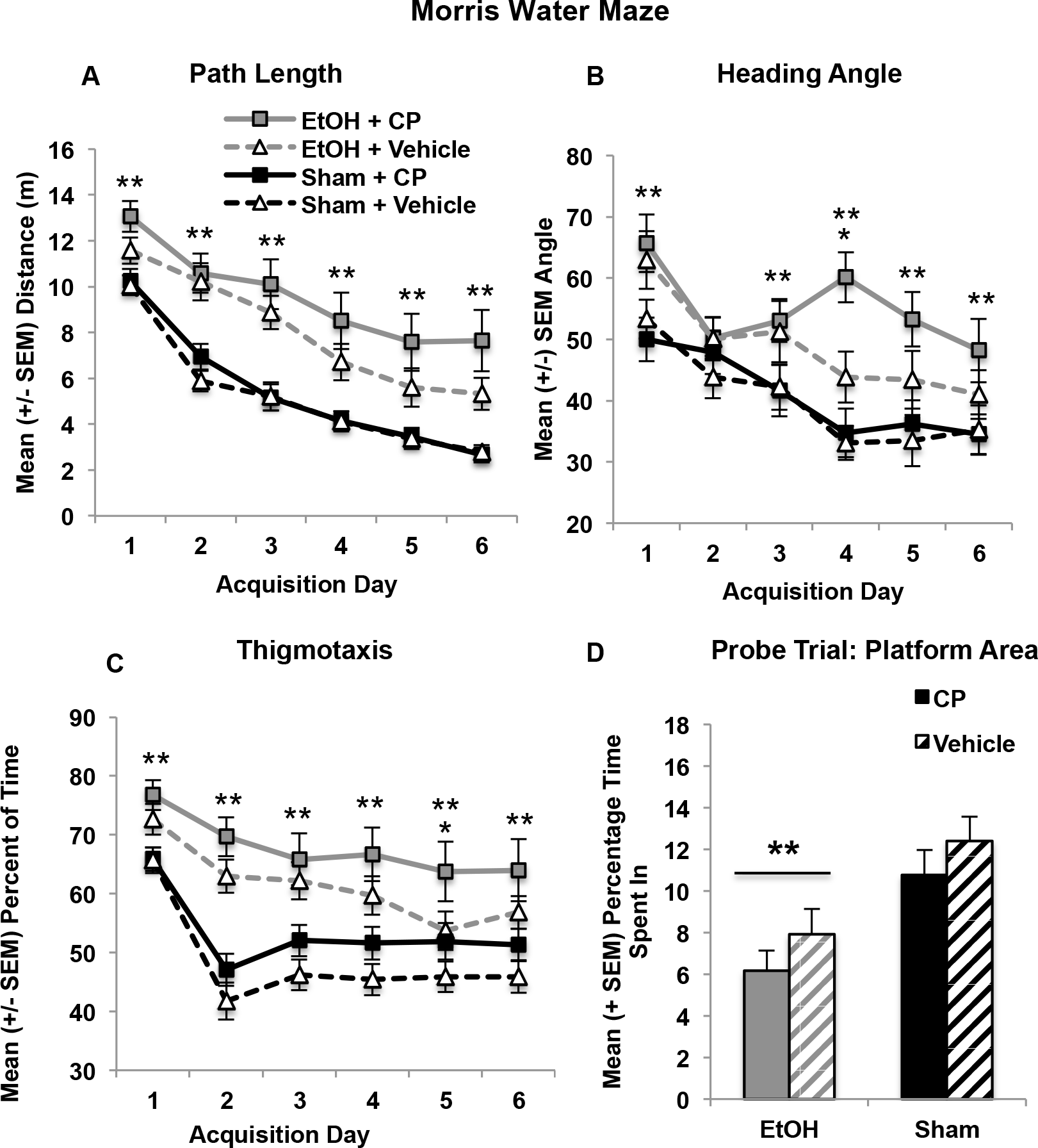
Developmental EtOH exposure impaired performance on the spatial learning task (A, B), whereas EtOH and CP separately increased thigmotaxis during acquisition (C). Only developmental EtOH exposure significantly impaired spatial memory during the probe test. (D). ** EtOH < Sham.

In contrast, both developmental EtOH and CP exposure separately increased thigmotaxis (wall-hugging, outer 10% of the diameter) during acquisition (Figure 6C), but did not interact. Subjects exposed to EtOH spent more time (F[1,103] = 26.23, *p* < 0.001) and did not decline in thigmotaxis as quickly as sham-intubated subjects, producing an interaction of Day*EtOH (F[5,515] = 5.92, *p* < 0.001). Similarly, subjects exposed to CP spent more time in thigmotaxis across days (F[1,103] = 4.20, *p* < 0.05). The same effects were seen in thigmotaxic path lengths (data not shown).

#### 3.4.2. Test (Probe)

The probe test occurred on a single day, so a 2 (EtOH) × 2 (CP) × 2 (Sex) ANOVA was utilized. Only developmental EtOH exposure impaired performance during the probe trial. EtOH-exposed subjects spent less time in the platform area (3 times the diameter of the platform; 1 of 8 possible locations; F[1,103] = 14.41, *p* < 0.001; Figure 6D); similar effects were seen in the ratio of time spent in the target area versus all other areas as well as time spent in the platform quadrant (1 of 4 quadrants; data not shown). In addition, EtOH-exposed subjects also passed through the platform area less often, including in comparison to passes through all other areas combined (target-pass ratio, data not shown). No main effects or interactions with developmental CP exposure were observed in any measure. Overall, females performed worse than males, regardless of exposure type (*p’*s < 0.01).

### 3.5. Brain Weights

Developmental EtOH exposure decreased total gross brain weights (F[1,103] = 11.88, *p* < 0.01; Figure 7A), but did not alter the total brain-body weight ratios. EtOH exposure also decreased the forebrain (F[1,103] = 79.17, *p* < 0.001; Figure 7B) and cerebellum (F[1,103] = 59.72, *p* < 0.001; Figure 7C) weights, as well as the forebrain-(F[1,103] = 17.86, *p* < 0.001) and cerebellum-body weight ratios (F[1,103] = 37.03, *p* < 0.001)(data not shown). Developmental CP exposure did not significantly alter gross weights of any brain region. Finally, male subjects had greater weights and ratios for total brain, forebrain, and cerebellum, compared to female subjects (*p’*s < 0.001).

**Figure 7.**
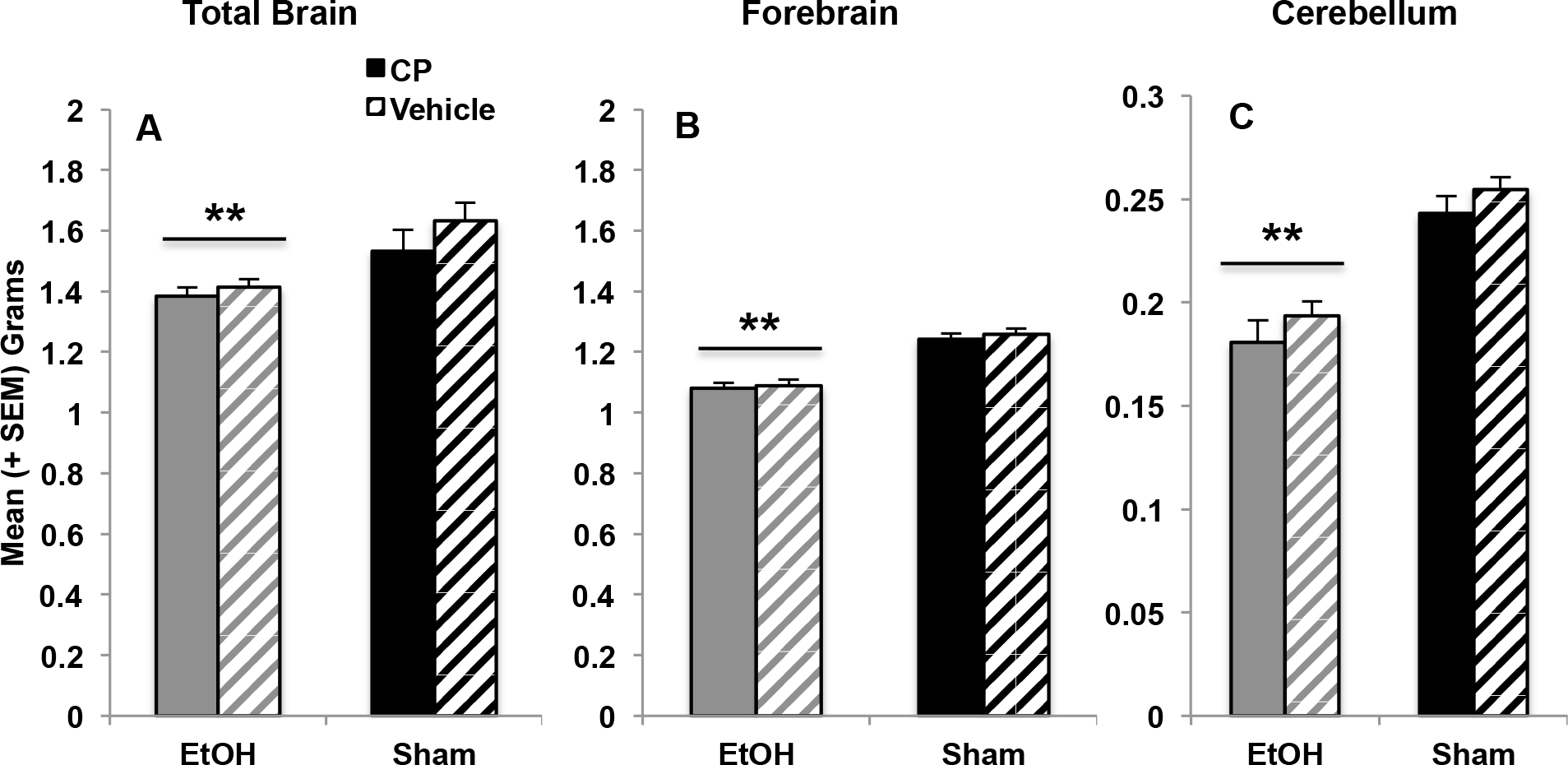
Developmental EtOH exposure reduced total brain weights (A) as well as forebrain (B) and cerebellum weights (C), whereas CP exposure had no long-lasting effects on brain weights.

## 4. Discussion

The current study suggests that exposure to combined alcohol and cannabinoids during early postnatal development may alter brain and behavioral development in a domain-specific manner. Importantly, concurrent exposure to alcohol and cannabinoids during development may disrupt behavioral development more than prenatal exposure to either drug alone. This was particularly evident in open field activity.

First, exposure to developmental alcohol increased overall activity levels, with elevations in locomotor activity, rearing, as well as entries and time spent in the center of the chamber. Consistent with the results of this study, past preclinical research has consistently shown that rats exposed to alcohol during early development display increased activity levels (Thomas et al., 2009; Thomas et al., 2007; Tran et al., 2000), which corresponds to clinical reports that children with FASD are commonly hyperactive (Mattson et al., 2011), although some clinical data fail to find difference in activity levels (Glass et al., 2014). Similarly, developmental CP exposure alone also increased locomotor activity, rearing and center travel and entries. Preclinical data examining activity levels following prenatal cannabis exposure are more limited and inconsistent. One study has shown that limited prenatal THC exposure (GD 10-12) increased activity levels from PD 9-21, yet did not increase time the spent within the center of the chamber (Borgen et al., 1973), which is consistent with the findings of the current study. However, other preclinical studies have shown no difference or even decreased activity levels in early adulthood (Fride et al., 1996). Limited clinical studies report that children (El Marroun et al., 2009; Hofman et al., 2004; Jaddoe et al., 2012; Jaddoe et al., 2010) and adolescents (Fried et al., 1988) prenatally exposed to marijuana exhibit hyperactivity, further indicating that exposure to cannabinoids may have long-lasting effects on activity levels.

The effects of combined developmental alcohol and THC were largely additive, with subjects exposed to both drugs exhibiting the most severe increases in activity levels. However, there were some synergistic effects, as subjects exposed to combined developmental alcohol and cannabinoids habituated more slowly compared to all other groups. This alteration may represent unique deficits in simple learning, as habituation represents learning of the environment. While these measures have not been well examined in preclinical studies, clinical reports do suggest that children (Goldschmidt et al., 2000) and adolescents (Fried et al., 1988) prenatally exposed to marijuana exhibit impairments in attention and impulsivity in addition to hyperactivity. It would be important to investigate the effects of combined alcohol and cannabinoids on habituation of other behaviors to better understand the nature of this behavioral change.

The effects of both drugs on emotional development are more complex to interpret. Increases in locomotion and time spent in the center of the open field chamber may indicate that the EtOH-exposed subjects are less anxious or are more inclined to take risks (Prut et al., 2003), suggesting that developmental alcohol exposure may affect emotional development as well as activity. However, developmental alcohol exposure did not alter the time spent in the open versus closed arms of the elevated plus maze. Ethanol exposure itself did decrease grooming behaviors, an ambiguous behavior as it has been interpreted as a representation of increased or decreased anxiety levels (Rodgers et al., 1997). In contrast, subjects exposed to CP did increase the time spent in the open arms in the elevated plus maze, suggesting that developmental cannabinoid exposure leads to reductions in anxiety. Yet CP exposure did not significantly affect time spent in the center of the open field. Importantly, the contexts of the open field and elevated plus maze vary considerably and subjects were tested at different ages. In sum, based on elevated plus maze, one could conclude that CP, but not EtOH, exposure during the 3^rd^ trimesters reduces anxiety, although given behaviors in the open field, one must be cautious of making firm conclusions of the effects of both drugs on stress and emotional development.

Moreover, both ethanol and CP, by themselves, increased thigmotaxis in the spatial learning task. The combination produced additive effects, as subjects exposed to both drugs were more thigmotaxic than subjects exposed to only one drug. Thigmotaxis has classically been described as increased fear or anxiety (Barnett, 1963), but more recent research suggests it may also represent a cognitive impairment of selection-response (Devan et al., 1999). Given the results of the open-field activity and elevated plus maze tasks, where drug-exposed subjects spend more time in open areas, the increased thigmotaxis among subjects could suggest a specific difference in cognitive strategy during the spatial learning task.

Interestingly, exposure to alcohol, and not CP, during the 3^rd^ trimester equivalent impaired spatial learning and memory, producing longer path lengths to find the platform and larger heading angles during acquisition, as well as impaired performance on the probe trial. This spatial learning deficit is consistent with past literature from our lab and others (Berman et al., 2000; Girard et al., 2000; Thomas et al., 2007). Subjects exposed to cannabinoids during development did not show these same impairments; this is not consistent with some past literature suggesting that prenatal THC can impair spatial learning and memory (O’shea et al., 2005), although a different task was used in that study. It is also somewhat surprising, given the high density of cannabinoid receptors in the hippocampus. Nevertheless, the current study found no evidence that early postnatal exposure to cannabinoids influence spatial memory, nor that cannabinoids exacerbate impairments related to developmental alcohol exposure.

Developmental alcohol exposure decreased body weights throughout behavioral testing. Cannabinoid exposure exacerbated these body weight reductions up through subjects’ early adolescent periods (PD 25), but did not have lasting effects. This is consistent with past clinical (Riley et al., 2011) and preclinical data (Ryan et al., 2008; Thomas et al., 2009; Thomas et al., 2004) showing that prenatal alcohol exposure can have long-lasting effects on body weight, whereas the effects of prenatal cannabis exposure on physical development have been described as subtle and/or brief in past clinical reviews (Day et al., 1991a; Day et al., 1991b; Fergusson et al., 2002; Fried et al., 1987; Huizink, 2014; Hurd et al., 2005). Similarly, developmental alcohol exposure led to long-lasting reductions in brain weight, whereas developmental cannabinoid exposure had no significant effects. Brain weights are a gross measure of neuropathology; however, they indicate how devastating early alcohol may be on development. Relatively little is known of the effects of prenatal cannabis exposure on more precise measures of neuropathology, but is desperately needed. Future research would benefit from detailed neuropathological analyses following combined prenatal alcohol and cannabinoid exposure, as past research has shown that the combination can be neurotoxic when either drug alone is not (Hansen et al., 2008).

The current study found limited overall sex differences in behavioral development, and these differences were independent of developmental alcohol or cannabinoid exposure.

Previous research has shown sex-dependent effects of developmental alcohol exposure can be found in spatial learning (Goodlett et al., 1995; Kelly et al., 1988; Zimmerberg et al., 1991), stress responsivity (Weinberg, 1992; Weinberg et al., 2008), and neuropathology (Barron et al., 1988; Zimmerberg et al., 1990; Zimmerberg et al., 1989), although the patterns have been inconsistent and often sex interactions are not seen (Thomas et al., 2009; Thomas et al., 2004; Thomas et al., 2000). Similarly, sex-dependent effects of prenatal cannabis exposure have also been reported on emotional behaviors, motor performance, and physiology (Biscaia et al., 2003; Navarro et al., 1994; Pérez-Rosado et al., 2000; Rubino et al., 2008). In general, these studies suggest that females may be more sensitive to the effects of prenatal drug exposure. Although females performed worse on the Morris water maze, regardless of exposure group, no other sex differences were observed in the present study.

There are some limitations to the present study. The dose of CP was chosen to reflect the amount of THC consumed by women of childbearing age in the general population (Mehmedic et al., 2010) and is based on doses used in other studies, including developmental (Gilbert et al., 2016) and adult (Gilbert et al., 2016; LaFleur et al., 2018; Maguire et al., 2016; Minervini et al., 2017) CP exposure. The dose of ethanol used in the current study is a well-established model of moderate/high binge-like third trimester exposure that historically produces robust behavioral deficits after exposure has ceased (Idrus et al., 2011; Klintsova et al., 2002; Ryan et al., 2008; Thomas et al., 2007; Thomas et al., 2000); the ethanol-related deficits seen in the current study are similar to those our laboratory has observed in the past. However, follow-up studies should determine how the combinations of various doses of ethanol and/or cannabinoids affect development, including sub-threshold doses of each drug. For example, combined exposure to cannabinoids and alcohol during early postnatal development has been shown to be synergistically neurotoxic, even when each drug is administered at a sub-threshold level (Hansen et al., 2008).

Importantly, although CP-55,940 is a cannabinoid receptor agonist similar to THC, it is not structurally chemically identical to THC. CP is more potent than THC, but the two substances have similar peak effects, durations of action, and neurobehavioral effects (McGregor et al., 1996). Given that cross-tolerance can develop between THC and CP, these substances have certain characteristics in common; however, it should be noted that there may be differences in specific mechanisms of action such as spontaneous activity, hypothermia, and catalepsy (Fan et al., 1994), which could be partially due to CP’s shorter half-life (Fouda et al., 1987; Grotenhermen, 2003). The CP dose used in this study is similar to a 12 mg/kg THC dose; i.p. injections of 10 mg/kg THC in a rat produce plasma levels around 65 ng/mL (Javadi-Paydar et al., 2018), which is consistent with human consumption levels (Desrosiers et al., 2014). Dose ranges in rat studies are highly variable, but recent studies have used THC doses generally ranging from 10-30 mg/kg when mimicking high potencies (Farquhar et al., 2019; Nguyen et al., 2016; Taffe et al., 2015; Wakley et al., 2014).

In addition, although we did not directly measure body temperature, we did videotape pup and maternal behavior and did not find any differences in behavior. In addition, during the drug exposures, pups were removed from the dam for only a few minutes during the drug exposure. When removed from the home cage during this time, they were maintained on a heating pad with littermates. The mortality rates, peak blood alcohol concentrations (BACs), and body weights of these subjects during the exposure period are explained in detail elsewhere (Breit et al., 2018, under revision). Briefly, CP did increase peak BACs, but only among females, one of the few sex differences we have found. CP by itself had no effect on mortality rates or body weights during exposure, although these measures were more affected by the combination of CP and ethanol.

Lastly, it is important to acknowledge that the behavioral testing utilized a within-subjects design, where all subjects performed each behavioral task, and each test occurred at different ages. However, the age of testing was determined based on the ontogeny of the behavior and the developmental ages at which these tasks are sensitive to early alcohol exposure, (Idrus et al., 2011; Osborn et al., 1998; Ryan et al., 2008) and the testing order was designed to minimize carryover effects. Nevertheless, the present data do indicate that cannabinoid exposure may disrupt behavioral development by itself and may exacerbate some behavioral alterations induced by developmental alcohol exposure.

## 5. Conclusions

In conclusion, the current study suggests that exposure to alcohol and cannabinoids during a period of development equivalent to late gestation may have individual, additive, and synergistic effects on behavioral development in domain-specific manners. Only developmental alcohol exposure impaired spatial learning and produced gross brain pathology. In contrast, only developmental cannabinoid exposure increased time spent in the open arms of the plus maze. However, both increased open field activity levels and increased thigmotaxis in the water maze, with largely additive consequences. Importantly, combined exposure to alcohol and cannabinoids may specifically impair simple learning processes, such as habituation, in a synergistic manner. These results show that the combination of ethanol and cannabinoids can produce more severe behavioral alterations than either drug alone, findings that have important implications for individuals exposed to both alcohol and marijuana during gestation, including an increased risk of behavioral problems later in life.

## Acknowledgements

Supported by NIAAA grant AA025425 to Dr. Thomas and an NIH Loan Repayment Program award to Dr. Breit. Special thanks to Dr. Nirelia Idrus, Cristina Rodriguez, and the Center for Behavioral Teratology at San Diego State University for assisting in data collection and interpretation.

## REFERENCES

Abel, E., & Subramanian, M. (1990). Effects of low doses of alcohol on delta-9-tetrahydrocannabinol’s effects in pregnant rats. Life Sciences, 47(18), 1677–1682.

Abel, E., Tan, S., & Subramanian, M. (1987). Effects of ∆9‐tetrahydrocannabinol, phenobarbital, and their combination on pregnancy and offspring in rats. Teratology, 36(2), 193–198.

Abel, E. L., Rockwood, G. A., & Riley, E. P. (1986). The effects of early marijuana exposure. Handbook of Behavioral Teratology (pp. 267–288): Springer.

Barnett, S. A. (1963). The rat: A study in behavior: Routledge.

Barron, S., Tiernan, S. B., & Riley, E. P. (1988). Effects of prenatal alcohol exposure on the sexually dimorphic nucleus of the preoptic area of the hypothalamus in male and female rats. Alcoholism: Clinical and Experimental Research, 12(1), 59–64.

Basavarajappa, B. S. (2015). Fetal alcohol spectrum disorder: Potential role of endocannabinoids signaling. Brain Sciences, 5(4), 456–493.

Basavarajappa, B. S., Ninan, I., & Arancio, O. (2008). Acute ethanol suppresses glutamatergic neurotransmission through endocannabinoids in hippocampal neurons. Journal of Neurochemistry, 107(4), 1001–1013.

Belue, R. C., Howlett, A. C., Westlake, T. M., & Hutchings, D. E. (1995). The ontogeny of cannabinoid receptors in the brain of postnatal and aging rats. Neurotoxicology and Teratology, 17(1), 25–30.

Berkovitz, R., Arieli, M., & Marom, E. (2011). Synthetic cannabinoids - the new” legal high” drugs. Harefuah, 150(12), 884–887, 937.

Berman, R. F., & Hannigan, J. H. (2000). Effects of prenatal alcohol exposure on the hippocampus: Spatial behavior, electrophysiology, and neuroanatomy. Hippocampus, 10(1), 94–110.

Berrendero, F., Sepe, N., Ramos, J. A., Di Marzo, V., & Fernández‐Ruiz, J. J. (1999). Analysis of cannabinoid receptor binding and mRNA expression and endogenous cannabinoid contents in the developing rat brain during late gestation and early postnatal period. Synapse, 33(3), 181–191.

Bhuvaneswar, C. G., Chang, G., Epstein, L. A., & Stern, T. A. (2007). Alcohol use during pregnancy: Prevalence and impact. Primary Care Companion to the Journal of Clinical Psychiatry, 9(6), 455–460.

Biscaia, M., Marín, S., Fernández, B., Marco, E. M., Rubio, M., Guaza, C., … Viveros, M. P. (2003). Chronic treatment with CP 55,940 during the peri-adolescent period differentially affects the behavioural responses of male and female rats in adulthood. Psychopharmacology, 170(3), 301–308. doi:10.1007/s00213-003-1550-7

Borgen, L. A., Davis, W. M., & Pace, H. B. (1973). Effects of prenatal ∆ 9-tetrahydrocannabinol on the development of rat offspring. Pharmacology Biochemistry and Behavior, 1(2), 203–206.

Botticelli, M. (2016). National drug control strategy: Data supplement 2015.

Botticelli, M. (2017). National drug control strategy: Data supplement 2016.

Breit, K. R., Zamudio, B., & Thomas, J. D. (2018). Altered motor development following late gestational alcohol and cannabinoid exposure in rats. Under review.

Centers for Disease Control and Prevention. (2015). Fetal alcohol spectrum disorders (FASD).

D’Alterio, S., & Bartke, A. (1979). Perinatal exposure to cannabinoids alters male reproductive function in mice. Science, 205(4413), 1420–1422.

Day, N., Sambamoorthi, U., Taylor, P., Richardson, G., Robles, N., Jhon, Y., … Jasperse, D. (1991a). Prenatal marijuana use and neonatal outcome. Neurotoxicology and Teratology, 13(3), 329–334.

Day, N. L., & Richardson, G. A. (1991b). Prenatal marijuana use: Epidemiology, methodologic issues, and infant outcome. Clinics in Perinatology, 18(1), 77–91.

de Salas-Quiroga, A., Díaz-Alonso, J., García-Rincón, D., Remmers, F., Vega, D., Gómez-Cañas, M., … Galve-Roperh, I. (2015). Prenatal exposure to cannabinoids evokes long-lasting functional alterations by targeting CB1 receptors on developing cortical neurons. Proceedings of the National Academy of Sciences, 112(44), 13693–13698.

Desrosiers, N. A., Himes, S. K., Scheidweiler, K. B., Concheiro-Guisan, M., Gorelick, D. A., & Huestis, M. A. (2014). Phase I and II cannabinoid disposition in blood and plasma of occasional and frequent smokers following controlled smoked cannabis. Clinical Chemistry, 60(4), 1–13.

Devan, B., McDonald, R., & White, N. (1999). Effects of medial and lateral caudate-putamen lesions on place-and cue-guided behaviors in the water maze: Relation to thigmotaxis. Behavioural Brain Research, 100(1-2), 5–14.

Dobbing, J., & Sands, J. (1979). Comparative aspects of the brain growth spurt. Early Human Development, 3(1), 79–83.

Ebrahim, S. H., & Gfroerer, J. (2003). Pregnancy-related substance use in the united states during 1996–1998. Obstetrics & Gynecology, 101(2), 374–379.

El Marroun, H., Hudziak, J. J., Tiemeier, H., Creemers, H., Steegers, E. A., Jaddoe, V. W., … Huizink, A. C. (2011). Intrauterine cannabis exposure leads to more aggressive behavior and attention problems in 18-month-old girls. Drug and Alcohol Dependence, 118(2-3), 470–474.

El Marroun, H., Tiemeier, H., Steegers, E. A., Jaddoe, V. W., Hofman, A., Verhulst, F. C., … Huizink, A. C. (2009). Intrauterine cannabis exposure affects fetal growth trajectories: The generation r study. Journal of the American Academy of Child & Adolescent Psychiatry, 48(12), 1173–1181.

Fan, F., Compton, D. R., Ward, S., Melvin, L., & Martin, B. R. (1994). Development of cross-tolerance between delta 9-tetrahydrocannabinol, CP 55,940 and WIN 55,212. Journal of Pharmacology and Experimental Therapeutics, 271(3), 1383–1390.

Farquhar, C. E., Breivogel, C. S., Gamage, T. F., Gay, E. A., Thomas, B. F., Craft, R. M., & Wiley, J. L. (2019). Sex, THC, and hormones: Effects on density and sensitivity of CB1 cannabinoid receptors in rats. Drug and Alcohol Dependence, 194, 20–27.

Fergusson, D. M., Horwood, L. J., & Northstone, K. (2002). Maternal use of cannabis and pregnancy outcome. BJOG: An International Journal of Obstetrics & Gynaecology, 109(1), 21–27.

Fernández-Ruiz, J., Berrendero, F., Hernández, M. L., & Ramos, J. A. (2000). The endogenous cannabinoid system and brain development. Trends in Neurosciences, 23(1), 14–20.

Fish, E. W., Boschen, K. E., Murdaugh, L. B., Mendoze-Romero, H. N., Williams, K., & Parnell, S. E. (2017). Alcohol exacerbates the teratogenic effects of prenatal cannabinoid exposure in a C57BL/6J mouse model. Birth Defects Research, 109(9), 688–688.

Fish, E. W., Gilbert, M. T., Sulik, K. K., & Parnell, S. E. (2016). Ethanol and the synthetic cannabinoid, CP-55,940, are synergistically teratogenic in a mouse model. Alcoholism: Clinical and Experimental Research, 40(S1), 287A–287A.

Fouda, H. G., Lukaszewicz, J., & Luther, E. W. (1987). Selected ion monitoring analysis of CP‐ 55,940, a cannabinoid derived analgetic agent. Biomedical & Environmental Mass Spectrometry, 14(11), 599–602.

Fride, E., & Mechoulam, R. (1996). Ontogenetic development of the response to anandamide and ∆9-tetrahydrocannabinol in mice. Developmental Brain Research, 95(1), 131–134.

Fried, P., & O’Connell, C. (1987). A comparison of the effects of prenatal exposure to tobacco, alcohol, cannabis and caffeine on birth size and subsequent growth. Neurotoxicology and Teratology, 9(2), 79–85.

Fried, P., & Watkinson, B. (1988). 12-and 24-month neurobehavioural follow-up of children prenatally exposed to marihuana, cigarettes and alcohol. Neurotoxicology and Teratology, 10(4), 305–313.

Fried, P. A., O’Connell, C. M., & Watkinson, B. (1992). 60-and 72-month follow-up of children prenatally exposed to marijuana, cigarettes, and alcohol: Cognitive and language assessment. Journal of Developmental & Behavioral Pediatrics, 13(6), 383–391.

Fried, P. A., & Watkinson, B. (1990). 36-and 48-month neurobehavioral follow-up of children prenatally exposed to marijuana, cigarettes, and alcohol. Journal of Developmental & Behavioral Pediatrics, 11(2), 49–58.

Garcı́a-Gil, L., Romero, J., Ramos, J. A., & Fernández-Ruiz, J. J. (1999). Cannabinoid receptor binding and mRNA levels in several brain regions of adult male and female rats perinatally exposed to ∆ 9-tetrahydrocannabinol. Drug and Alcohol Dependence, 55(1), 127–136.

Gauda, E. B. (2006). Knowledge gained from animal studies of the fetus and newborn: Application to the human premature infant. ILAR journal, 47(1), 1–4.

Gilbert, M. T., Sulik, K. K., Fish, E. W., Baker, L. K., Dehart, D. B., & Parnell, S. E. (2016). Dose-dependent teratogenicity of the synthetic cannabinoid CP-55,940 in mice. Neurotoxicology and Teratology, 58, 15–22.

Girard, T., Xing, H. C., Ward, G., & Wainwright, P. (2000). Early postnatal ethanol exposure has long‐term effects on the performance of male rats in a delayed matching‐to‐place task in the morris water maze. Alcoholism: Clinical and Experimental Research, 24(3), 300–306.

Glass, L., Graham, D. M., Deweese, B. N., Jones, K. L., Riley, E. P., & Mattson, S. N. (2014). Correspondence of parent report and laboratory measures of inattention and hyperactivity in children with heavy prenatal alcohol exposure. Neurotoxicology and Teratology, 42, 43–50.

Goldschmidt, L., Day, N. L., & Richardson, G. A. (2000). Effects of prenatal marijuana exposure on child behavior problems at age 10. Neurotoxicology and Teratology, 22(3), 325–336.

Goldschmidt, L., Richardson, G. A., Cornelius, M. D., & Day, N. L. (2004). Prenatal marijuana and alcohol exposure and academic achievement at age 10. Neurotoxicology and Teratology, 26(4), 521–532.

Gómez, M., Hernández, M., Johansson, B., De Miguel, R., Ramos, J. A., & Fernández-Ruiz, J. (2003). Prenatal cannabinoid exposure and gene expression for neural adhesion molecule L1 in the fetal rat brain. Developmental Brain Research, 147(1), 201–207.

Goodlett, C. R., & Johnson, T. B. (1997). Neonatal binge ethanol exposure using intubation: Timing and dose effects on place learning. Neurotoxicology and Teratology, 19(6), 435–446.

Goodlett, C. R., & Peterson, S. D. (1995). Sex differences in vulnerability to developmental spatial learning deficits induced by limited binge alcohol exposure in neonatal rats. Neurobiology of Learning and Memory, 64(3), 265–275.

Goodlett, C. R., Thomas, J. D., & West, J. R. (1991). Long-term deficits in cerebellar growth and rotarod performance of rats following “binge-like” alcohol exposure during the neonatal brain growth spurt. Neurotoxicology and Teratology, 13(1), 69–74.

Grotenhermen, F. (2003). Pharmacokinetics and pharmacodynamics of cannabinoids. Clinical Pharmacokinetics, 42(4), 327–360.

Hansen, H. H., Krutz, B., Sifringer, M., Stefovska, V., Bittigau, P., Pragst, F., … Ikonomidou, C. (2008). Cannabinoids enhance susceptibility of immature brain to ethanol neurotoxicity. Annals of Neurology, 64(1), 42–52.

Hofman, A., Jaddoe, V. W., Mackenbach, J. P., Moll, H. A., Snijders, R. F., Steegers, E. A., … Büller, H. A. (2004). Growth, development and health from early fetal life until young adulthood: The generation r study. Paediatric and Perinatal Epidemiology, 18(1), 61–72.

Huizink, A. (2014). Prenatal cannabis exposure and infant outcomes: Overview of studies. Progress in Neuro-Psychopharmacology and Biological Psychiatry, 52, 45–52.

Huizink, A. C., & Mulder, E. J. (2006). Maternal smoking, drinking or cannabis use during pregnancy and neurobehavioral and cognitive functioning in human offspring. Neuroscience & Biobehavioral Reviews, 30(1), 24–41.

Hurd, Y., Wang, X., Anderson, V., Beck, O., Minkoff, H., & Dow-Edwards, D. (2005). Marijuana impairs growth in mid-gestation fetuses. Neurotoxicology and Teratology, 27(2), 221–229.

Idrus, N. M., McGough, N. N. H., Spinetta, M. J., Thomas, J. D., & Riley, E. P. (2011). The effects of a single memantine treatment on behavioral alterations associated with binge alcohol exposure in neonatal rats. Neurotoxicology and Teratology, 33(4), 444–450.

Jaddoe, V. W., Van Duijn, C. M., Franco, O. H., Van Der Heijden, A. J., Van IIzendoorn, M. H., De Jongste, J. C., … Raat, H. (2012). The generation r study: Design and cohort update 2012. European Journal of Epidemiology, 27(9), 739–756.

Jaddoe, V. W., van Duijn, C. M., van der Heijden, A. J., Mackenbach, J. P., Moll, H. A., Steegers, E. A., … Hofman, A. (2010). The generation r study: Design and cohort update 2010. European Journal of Epidemiology, 25(11), 823–841.

Javadi-Paydar, M., Nguyen, J. D., Kerr, T. M., Grant, Y., Vandewater, S. A., Cole, M., & Taffe, M. A. (2018). Effects of ∆9-THC and cannabidiol vapor inhalation in male and female rats. Psychopharmacology, 9, 2541–2557.

Kelly, S. J., Goodlett, C. R., Hulsether, S. A., & West, J. R. (1988). Impaired spatial navigation in adult female but not adult male rats exposed to alcohol during the brain growth spurt. Behavioural Brain Research, 27(3), 247–257.

Khoury, J. E., Milligan, K., & Girard, T. A. (2015). Executive functioning in children and adolescents prenatally exposed to alcohol: A meta-analytic review. Neuropsychology Review, 25(2), 149–170.

Klintsova, A. Y., Scamra, C., Hoffman, M., Napper, R. M., Goodlett, C. R., & Greenough, W. T. (2002). Therapeutic effects of complex motor training on motor performance deficits induced by neonatal binge-like alcohol exposure in rats:: II. A quantitative stereological study of synaptic plasticity in female rat cerebellum. Brain Research, 937(1-2), 83–93.

LaFleur, R. A., Wilson, R. P., Morgan, D. J., & Henderson-Redmond, A. N. (2018). Sex differences in antinociceptive response to ∆-9-tetrahydrocannabinol and CP 55,940 in the mouse formalin test. NeuroReport, 29(6), 447–452.

Lange, S., Shield, K., Koren, G., Rehm, J., & Popova, S. (2014). A comparison of the prevalence of prenatal alcohol exposure obtained via maternal self-reports versus meconium testing: A systematic literature review and meta-analysis. BMC Pregnancy and Childbirth, 14(1), 127–138.

Lendoiro, E., González-Colmenero, E., Concheiro-Guisán, A., de Castro, A., Cruz, A., López-Rivadulla, M., & Concheiro, M. (2013). Maternal hair analysis for the detection of illicit drugs, medicines, and alcohol exposure during pregnancy. Therapeutic Drug Monitoring, 35(3), 296–304.

Little, P., Compton, D., Johnson, M., Melvin, L., & Martin, B. (1988). Pharmacology and stereoselectivity of structurally novel cannabinoids in mice. Journal of Pharmacology and Experimental Therapeutics, 247(3), 1046–1051.

Maguire, D. R., & France, C. P. (2016). Additive antinociceptive effects of mixtures of the kappa opioid receptor agonist spiradoline and the cannabinoid receptor agonist CP55940 in rats. Behavioural Pharmacology, 27(1), 69.

Mattson, S. N., Crocker, N., & Nguyen, T. T. (2011). Fetal Alcohol Spectrum Disorders: Neuropsychological and Behavioral Features. Neuropsychology Review, 21(2), 81–101. doi:10.1007/s11065-011-9167-9

McGough, N. N., Thomas, J. D., Dominguez, H. D., & Riley, E. P. (2009). Insulin-like growth factor-I mitigates motor coordination deficits associated with neonatal alcohol exposure in rats. Neurotoxicology and Teratology, 31(1), 40–48.

McGregor, I. S., Issakidis, C. N., & Prior, G. (1996). Aversive effects of the synthetic cannabinoid CP-55,940 in rats. Pharmacology Biochemistry and Behavior, 53(3), 657–664.

Mehmedic, Z., Chandra, S., Slade, D., Denham, H., Foster, S., Patel, A. S., … ElSohly, M. A. (2010). Potency trends of ∆9THC and other cannabinoids in confiscated cannabis preparations from 1993 to 2008. Journal of Forensic Sciences, 55(5), 1209–1217.

Mereu, G., Fà, M., Ferraro, L., Cagiano, R., Antonelli, T., Tattoli, M., … Cuomo, V. (2003). Prenatal exposure to a cannabinoid agonist produces memory deficits linked to dysfunction in hippocampal long-term potentiation and glutamate release. Proceedings of the National Academy of Sciences, 100(8), 4915–4920.

Midanik, L. T. (1988). Validity of self‐reported alcohol use: A literature review and assessment. British Journal of Addiction, 83(9), 1019–1029.

Minervini, V., Dahal, S., & France, C. P. (2017). Behavioral characterization of κ opioid receptor agonist spiradoline and cannabinoid receptor agonist CP55940 mixtures in rats. Journal of Pharmacology and Experimental Therapeutics, 360(2), 280–287.

Moore, E. M., Migliorini, R., Infante, M. A., & Riley, E. P. (2014). Fetal alcohol spectrum disorders: Recent neuroimaging findings. Current Developmental Disorders Reports, 1(3), 161–172.

Nagre, N. N., Subbanna, S., Shivakumar, M., Psychoyos, D., & Basavarajappa, B. S. (2015). CB1-receptor knockout neonatal mice are protected against ethanol-induced impairments of DNMT 1, DNMT 3A, and DNA methylation. Journal of Neurochemistry, 132(4), 429–442.

Navarro, M., Rubio, P., & Rodríguez de Fonseca, F. (1994). Sex-dimorphic psychomotor activation after perinatal exposure to (−)-∆9-tetrahydrocannabinol. An ontogenic study in wistar rats. Psychopharmacology, 116(4), 414–422. doi:10.1007/bf02247471

Nguyen, J. D., Aarde, S. M., Vandewater, S. A., Grant, Y., Stouffer, D. G., Parsons, L. H., … Taffe, M. A. (2016). Inhaled delivery of ∆9-tetrahydrocannabinol (THC) to rats by e-cigarette vapor technology. Neuropharmacology, 109, 112–120.

Norman, A. L., O’brien, J. W., Spadoni, A. D., Tapert, S. F., Jones, K. L., Riley, E. P., & Mattson, S. N. (2013). A functional magnetic resonance imaging study of spatial working memory in children with prenatal alcohol exposure: Contribution of familial history of alcohol use disorders. Alcoholism: Clinical and Experimental Research, 37(1), 132–140.

O’shea, M., & Mallet, P. (2005). Impaired learning in adulthood following neonatal ∆9-THC exposure. Behavioural Pharmacology, 16(5-6), 455–461.

Olney, J. W., Wozniak, D. F., Jevtovic-Todorovic, V., Farber, N. B., Bittigau, P., & Ikonomidou, C. (2002). Drug-induced apoptotic neurodegeneration in the developing brain. Brain Pathology, 12(4), 488–498.

Osborn, J., Kim, C., Steiger, J., & Weinberg, J. (1998). Prenatal ethanol exposure differentially alters behavior in males and females on the elevated plus maze. Alcoholism: Clinical and Experimental Research, 22(3), 685–696.

Pérez-Rosado, A., Manzanares, J., Fernández-Ruiz, J., & Ramos, J. A. (2000). Prenatal ∆9-tetrahydrocannabinol exposure modifies proenkephalin gene expression in the fetal rat brain: Sex-dependent differences. Developmental Brain Research, 120(1), 77–81.

Prut, L., & Belzung, C. (2003). The open field as a paradigm to measure the effects of drugs on anxiety-like behaviors: A review. European Journal of Pharmacology, 463(1-3), 3–33.

Richardson, G. A., Ryan, C., Willford, J., Day, N. L., & Goldschmidt, L. (2002). Prenatal alcohol and marijuana exposure: Effects on neuropsychological outcomes at 10 years. Neurotoxicology and Teratology, 24(3), 309–320.

Riley, E. P., Infante, M. A., & Warren, K. R. (2011). Fetal alcohol spectrum disorders: An overview. Neuropsychology Review, 21(2), 73–80. doi:10.1007/s11065-011-9166-x

Rodgers, R., & Dalvi, A. (1997). Anxiety, defence and the elevated plus-maze. Neuroscience & Biobehavioral Reviews, 21(6), 801–810.

Rodriguez de Fonseca, F., Ramos, J. A., Bonnin, A., & Fernández-Ruiz, J. J. (1993). Presence of cannabinoid binding sites in the brain from early postnatal ages. Neuroreport, 4(2), 135–138.

Roozen, S., Peters, G. J. Y., Kok, G., Townend, D., Nijhuis, J., & Curfs, L. (2016). Worldwide prevalence of fetal alcohol spectrum disorders: A systematic literature review including meta‐analysis. Alcoholism: Clinical and Experimental Research, 40(1), 18–32.

Rubino, T., Realini, N., Guidali, C., Braida, D., Capurro, V., Castiglioni, C., … Sala, M. (2008). Chronic ∆ 9-tetrahydrocannabinol during adolescence provokes sex-dependent changes in the emotional profile in adult rats: Behavioral and biochemical correlates. Neuropsychopharmacology, 33(11), 2760–2771.

Ryan, S. H., Williams, J. K., & Thomas, J. D. (2008). Choline supplementation attenuates learning deficits associated with neonatal alcohol exposure in the rat: Effects of varying the timing of choline administration. Brain Research, 1237, 91–100.

Saint Louis, C. (2017). Pregnant women turn to marijuana, perhaps harming infants. The New York Times. Retrieved from https://www.nytimes.com/2017/02/02/health/marijuana-and-pregnancy.html?emc=eta1&_r=0

Schneider, M. (2009). Cannabis use in pregnancy and early life and its consequences: Animal models. European Archives of Psychiatry and Clinical Neuroscience, 259(7), 383–393.

Schneider, M. L., Moore, C. F., & Adkins, M. M. (2011). The effects of prenatal alcohol exposure on behavior: Rodent and primate studies. Neuropsychology Review, 21(2), 186–203.

Smith, A. M., Fried, P. A., Hogan, M. J., & Cameron, I. (2006). Effects of prenatal marijuana on visuospatial working memory: An fMRI study in young adults. Neurotoxicology and Teratology, 28(2), 286–295.

Subbanna, S., Shivakumar, M., Psychoyos, D., Xie, S., & Basavarajappa, B. S. (2013). Anandamide–CB1 receptor signaling contributes to postnatal ethanol-induced neonatal neurodegeneration, adult synaptic, and memory deficits. Journal of Neuroscience, 33(15), 6350–6366.

Subbaraman, M. S., & Kerr, W. C. (2015). Simultaneous versus concurrent use of alcohol and cannabis in the national alcohol survey. Alcoholism: Clinical and Experimental Research, 39(5), 872–879.

Substance Abuse and Mental Health Services Administration. (2015). Behavioral Health Trends in the United States: Results from the 2014 National Survey on Drug Use and Health.

Substance Use and Mental Health Services Administration. (2014). Results from the 2013 National Survey on Drug Use and Health: Summary of National Findings.

Taffe, M. A., Creehan, K. M., & Vandewater, S. A. (2015). Cannabidiol fails to reverse hypothermia or locomotor suppression induced by ∆9‐tetrahydrocannabinol in Sprague‐Dawley rats. British journal of pharmacology, 172(7), 1783–1791.

Tai, S., & Fantegrossi, W. E. (2014). Synthetic cannabinoids: Pharmacology, behavioral effects, and abuse potential. Current Addiction Reports, 1(2), 129–136.

Taylor, P. A., Jacobson, S. W., van der Kouwe, A., Molteno, C. D., Chen, G., Wintermark, P., … Meintjes, E. M. (2015). A DTI‐based tractography study of effects on brain structure associated with prenatal alcohol exposure in newborns. Human Brain Mapping, 36(1), 170–186.

Thomas, J. D., Abou, E. J., & Dominguez, H. D. (2009). Prenatal choline supplementation mitigates the adverse effects of prenatal alcohol exposure on development in rats. Neurotoxicology and Teratology, 31(5), 303–311.

Thomas, J. D., Biane, J. S., O’bryan, K. A., O’neill, T. M., & Dominguez, H. D. (2007). Choline supplementation following third-trimester-equivalent alcohol exposure attenuates behavioral alterations in rats. Behavioral Neuroscience, 121(1), 120–130.

Thomas, J. D., Garrison, M., & O’Neill, T. M. (2004). Perinatal choline supplementation attenuates behavioral alterations associated with neonatal alcohol exposure in rats. Neurotoxicology and Teratology, 26(1), 35–45.

Thomas, J. D., La Fiette, M. H., Quinn, V. R., & Riley, E. P. (2000). Neonatal choline supplementation ameliorates the effects of prenatal alcohol exposure on a discrimination learning task in rats. Neurotoxicology and Teratology, 22(5), 703–711.

Thomas, J. D., Sather, T. M., & Whinery, L. A. (2008). Voluntary exercise influences behavioral development in rats exposed to alcohol during the neonatal brain growth spurt. Behavioral Neuroscience, 122(6), 1264–1273.

Thomas, J. D., & Tran, T. D. (2012). Choline supplementation mitigates trace, but not delay, eyeblink conditioning deficits in rats exposed to alcohol during development. Hippocampus, 22(3), 619–630.

Tran, T. D., Cronise, K., Marino, M. D., Jenkins, W. J., & Kelly, S. J. (2000). Critical periods for the effects of alcohol exposure on brain weight, body weight, activity and investigation. Behavioural Brain Research, 116(1), 99–110.

Trezza, V., Campolongo, P., Manduca, A., Morena, M., Palmery, M., Vanderschuren, L., & Cuomo, V. (2012). Altering endocannabinoid neurotransmission at critical developmental ages: Impact on rodent emotionality and cognitive performance. Frontiers in Behavioral Neuroscience, 6, 1–12.

Uban, K., Herting, M., Wozniak, J., Sowell, E., & CIFASD. (2017). Sex differences in associations between white matter microstructure and gonadal hormones in children and adolescents with prenatal alcohol exposure. Psychoneuroendocrinology, 83, 111–121.

Wakley, A. A., Wiley, J. L., & Craft, R. M. (2014). Sex differences in antinociceptive tolerance to delta-9-tetrahydrocannabinol in the rat. Drug and Alcohol Dependence, 143, 22–28.

Weinberg, J. (1992). Prenatal ethanol effects: Sex differences in offspring stress responsiveness. Alcohol, 9(3), 219–223.

Weinberg, J., Sliwowska, J. H., Lan, N., & Hellemans, K. (2008). Prenatal alcohol exposure: Foetal programming, the hypothalamic‐pituitary‐adrenal axis and sex differences in outcome. Journal of Neuroendocrinology, 20(4), 470–488.

West, J. R., Hamre, K. M., & Pierce, D. R. (1984). Delay in brain growth induced by alcohol in artificially reared rat pups. Alcohol, 1(3), 213–222.

Zimmerberg, B., & Mickus, L. A. (1990). Sex differences in corpus callosum: Influence of prenatal alcohol exposure and maternal undernutrition. Brain Research, 537(1), 115–122.

Zimmerberg, B., & Scalzi, L. V. (1989). Commissural size in neonatal rats: Effects of sex and prenatal alcohol exposure. International Journal of Developmental Neuroscience, 7(1), 81–86.

Zimmerberg, B., Sukel, H. L., & Stekler, J. D. (1991). Spatial learning of adult rats with fetal alcohol exposure: Deficits are sex-dependent. Behavioural Brain Research, 42(1), 49–56.

